# Genomic characterization of plasmids harboring *bla*_NDM-1,-5,-7_ carbapenemase alleles in clinical *Klebsiella pneumoniae* in Pakistan

**DOI:** 10.1101/2024.09.06.611696

**Authors:** Muhammad Usman Qamar, Roberto Sierra, Kokab Jabeen, Muhammad Rizwan, Ayesha Rashid, Yumna Fatima Dar, Diego O. Andrey

**Affiliations:** Department of Microbiology and Molecular Medicine, Faculty of Medicine, University of Geneva, Switzerland; Institute of Microbiology, Faculty of Life Sciences, Government College University Faisalabad, 38000, Pakistan; Infectious Diseases Division, Geneva University Hospitals, Switzerland; Division of Laboratory Medicine, Geneva University Hospitals, Switzerland; Ameer ud Din Medical College/Lahore General Hospital and Postgraduate Medical Institute, Lahore, Pakistan

**Keywords:** CR-KP, plasmid, carbapenemases, NDM, antibiotic resistance, mobile genetic elements

## Abstract

*Klebsiella pneumoniae* is notorious for causing healthcare-associated infections, which become more complicated by the acquisition of *bla*_NDM_ genes via mobile genetic elements. Although Pakistan is a well-established hot spot of *bla*_NDM_-positive *K. pneumoniae*, detailed molecular descriptions of *bla*_NDM_-carrying plasmids are scarce. Seven *K. pneumoniae* isolates harboring *bla*_NDM_ were recovered from clinical sample sources during a six-month period and tested for antimicrobial susceptibility. A long-read approach was used for whole genome sequencing to obtain circularized plasmids and chromosomes for typing, annotation, and comparative analysis. The isolates were susceptible to colistin and tigecycline only among the tested antibiotics. We identified five STs: ST11, ST16, ST716, ST464, and ST2856. Notably, three strains possessed the hypervirulent capsule KL2, while five were classified as O locus type O2a. Evidence of genetic diversity was further highlighted by the presence of four IncC plasmids harboring *bla*_NDM-1_, two IncX3 plasmids harboring *bla*_NDM-5_, and a single hybrid IncFIB/IncHI1B plasmid harboring *bla*_NDM-7_. These plasmids also carried additional ARGs conferring resistance to aminoglycosides, cephalosporins, and fluoroquinolones. We identified the plasmidome of the *K. pneumoniae* isolates and characterized the NDM-carrying plasmids. Genetic analysis confirmed the presence of *bla*_NDM-1_ and *bla*_NDM-5_ on broad host range plasmids and *bla*_NDM-7_ in a previously unreported hybrid plasmid backbone. We emphasized the critical role of plasmids in spreading *bla*_NDM_ in the clinical setting in Pakistan. Hence, we stressed the urgent need for enhanced surveillance, not least in LMICs, infection control measures, and adherence to the AWaRe guidelines in antibiotics use.

## INTRODUCTION

Carbapenem-resistant *Klebsiella pneumoniae* (CR-KP) poses a grave global public health threat, particularly in low- and middle-income countries (LMICs), associated with significantly higher mortality rates (1, 2). This opportunistic pathogen may cause pneumonia, sepsis, urinary tract infections, and meningitis (3). CR-KP is categorized into the critical group by the World Health Organization (WHO) and is part of the ESKAPE pathogens, well-known for their role in healthcare-associated infections (4, 5). The Class B1 carbapenemase NDM was first identified on a 180 kb IncC plasmid in a clinical strain of *K. pneumoniae* (6), and over 60 NDM variants have been found in the β-lactamase database to date (7). *K. pneumoniae* accounted for more than half of NDM-positive Enterobacterales infections worldwide, followed by *Escherichia coli* and the *Enterobacter cloacae* complex (8, 9). Plasmids, as carriers of antimicrobial resistance genes (ARGs), play a pivotal role in acquiring and disseminating resistance. This is particularly evident in the case of CR-KP, which has expanded globally due to carbapenemase genes on mobile genetic elements (MGEs) such as plasmids, transposons, and insertion sequences via horizontal gene transfer (10). The Tn*125* transposons played a crucial role in disseminating the *bla*_NDM-1_ gene; however, in recent years, other elements, such as IS*26* and Tn*3000*, have exceeded their prevalence (11). The composite transposon Tn*125* is formed when two copies of IS*Aba125* capture a *bla*_NDM_ located on a plasmid or chromosome, facilitating the spread of *bla*_NDM_ to Enterobacterales and non-Enterobacterales (12). Notably, the bleomycin resistance gene (*ble*_MBL_) appears downstream of *bla*_NDM_, while IS*Aba125* is always found upstream and drives *ble*_MBL_ and *bla*_NDM_ co-expression (13). Recent data showed that *bla*_NDM_ genes are associated with at least 33 different types of plasmids, including IncC, IncX3, IncFIB, IncFII, IncH, and IncL/M, as well as untyped plasmids (11). *bla*_NDM-1_ and *bla*_NDM-5_ are frequently found on IncC and IncX3 plasmids, respectively, in Enterobacterales and have been reported from France (14), Italy (15), and China (16). In contrast, *bla*_NDM-7_ is less common and has been identified mainly on IncX3 plasmids, mostly in the *E. coli* (17). A few epidemiological studies from Pakistan indicated that the *bla*_NDM-1_ is present on different plasmid backbones in *K. pneumoniae*, but scant information is available on other alleles (18, 19). Our previous study was the first to report *bla*_NDM-5_ and *bla*_NDM-7_ in *K. pneumoniae* clinical isolates in Pakistan. These genes were located on plasmids ranging from 100kb to 150kb. However, genetic context analysis was not included (20). Therefore, in the present study, we characterized seven cases of NDM-producing *K. pneumoniae* recovered from clinical samples during a 6-month surveillance period (April to September 2023), and reconstructed the detailed genetic context of NDM-carrying plasmids.

## RESULTS

### Antimicrobial susceptibility (AST) of clinical isolates

Resistance to most medically important antibiotics, including β-lactam inhibitors (100%), cephalosporins (100%), carbapenems (100%), aminoglycosides (100%), fluoroquinolones (85.5%), and tetracycline (85.5%), was observed in these seven isolates. In Table 1, we show the antibiotic resistance profiles according to the WHO categories: “Access,” “Watch,” and “Reserve” (AWaRe). The isolates were resistant to most of the “Access” and “Watch” classes of antibiotics. However, no resistance was observed to colistin and tigecycline in the “Reserve” category. The bacterial multidrug resistance was assessed by calculating the Multiple Antibiotic Resistant (MAR) index using the formula a/b, where “a” represents the number of resistant antibiotics and “b” means the total number of antibiotics tested against the isolate. A threshold of ≤0.2 for the MAR index is generally used to indicate low-level resistance. The isolates in this investigation were deemed to possess a high degree of drug resistance. The MAR index varied between 0.73 and 0.86, as shown in Table 1.

**Table 1.** Antimicrobial susceptibility profile of CR-KP.

| WHO<br>AWaRe | Antibiotics<br>classification | Antibiotics<br>(ATC code) | MIC breakpoint<br>(µg/ml) | KPP-1 | KPP-2 | KPP-4 | KPP-5 | KPP-7 | KPP-9 | KPP-10 |
| --- | --- | --- | --- | --- | --- | --- | --- | --- | --- | --- |
| Access | β-lactamase inhibitor | AMC (J01CR02) | ≤8/4-≥32/16 | R | R | R | R | R | R | R |
|  | Co-trimoxazole | SXT (J01EE01) | ≤2/38-≥4/76 | R | R | R | S | R | R | R |
|  | Aminoglycoside | AK (J01GB06) | ≤4-≥16 | R | R | R | R | R | R | R |
|  | Tetracycline | DO (J01AA02) | ≤4-≥16 | R | R | R | R | R | R | S |
|  |  | TE (J01AA07) | ≤4-≥16 | R | R | S | R | R | R | R |
| Watch | Cephalosporins | CTX (J01DD01) | ≤1-≥4 | R | R | R | R | R | R | R |
|  |  | CRO (J01DD04) | ≤1-≥4 | R | R | R | R | R | R | R |
|  |  | CAZ (J01DD02) | ≤4-≥16 | R | R | R | R | R | R | R |
|  |  | FEP (J01DE01) | ≤2-≥16 | R | R | R | R | R | R | R |
|  | Carbapenems | IPM (J01DH51) | ≤1-≥4 | R | R | R | R | R | R | R |
|  |  | MEM (J01DH02) | ≤1-≥4 | R | R | R | R | R | R | R |
|  | Fluoroquinolones | CIP (J01MA02) | ≤0.25-≥1 | R | R | S | R | R | R | R |
|  |  | LEV (J01MA12) | ≤0.5-≥2 | R | R | R | R | R | R | R |
| Reserve | Polymyxins | CT (J01XB01) | ≥4 | S | S | S | S | S | S | S |
|  | Glycylcycline | TGC (J01AA12) | ≥2 | S | S | S | S | S | S | S |
| MAR Index | - | - | - | 0.86 | 0.86 | 0.73 | 0.80 | 0.86 | 0.86 | 0.80 |

AMC: amoxicillin/clavulanic acid, SXT: sulfamethoxazole/trimethoprim, AK: amikacin, DO: doxycycline, TE: tetracycline, CTX: cefotaxime, CRO: ceftriaxone, CAZ: ceftazidime, FEP: cefepime, IMP: imipenem, MEM: meropenem, CIP: ciprofloxacin, LEV: levofloxacin, CT: colistin, TGC: tigecycline. MAR index: multiple antibiotic index. ATC: Anatomical Therapeutic Chemical.

### Characterization of STs, capsular types, and ARGs

The MLST analysis of the seven CR-KP isolates revealed five distinct STs. Two isolates belonged to the high-risk clone ST11, two to ST716, and the remaining isolates belonged to ST464, ST2856, and ST16. These STs were mainly recovered from pus (n = 3) and blood samples (n = 2). The capsule (KL) and lipopolysaccharide O-antigen (O) locus types were utilized for typing *K. pneumoniae*. Three strains, two ST11, and one ST2856, were hypervirulent (hv) KL2, while two ST11, two ST716, and one ST2856 exhibited the O2a serotype. The ARGs analysis revealed three different NDM genotypes, including four *bla*_NDM-1_, two *bla*_NDM-5_, and one *bla*_NDM-7_. The seven isolates encoded a plethora of ARGs; the isolates KPP-1 and KPP-6 harbored 20 ARGs belonging to 12 different classes of antibiotics, followed by KPP-7 with 18, KPP-4 with 16, and KPP-2 with 15. The isolates co-harbored ARGs of carbapenems (*bla*_NDM_), cephalosporins (*bla*_TEM-1_, and *bla*_SHV-27_, *bla*_SHV-182_), aminoglycosides (*aac(3)-IId, rmtC, aac(6’)-Ib3, rmtB, aadA1*), fluoroquinolones (*qnr*S1, *qnr*B1), tetracycline (*tet*A, *tet*B), sulfonamides (*sul*1, *sul2)*, trimethoprim (*dfrA1, dfrA14, drfA27*), rifampicin (*ARR-3*), fosfomycin (*fosA, fosA5*), and chloramphenicol (*catB3*) (Figure 1).

**Figure 1.**
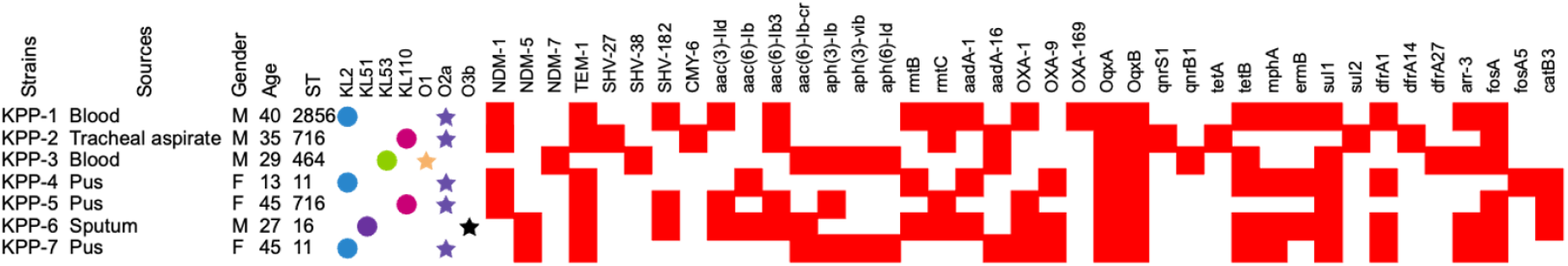
Source of specimens, patients’ demographics, clinical information, characterization of STs, capsule typing, and ARGs. Blue, green, and red circles indicate the K-locus, yellow, purple, and black stars indicate O-locus and red squares are the ARGs.

### Comparative analysis of *bla*_NDM_ encoding plasmids

NDM-1, -5, and -7 were encoded on IncC, IncX3, and hybrid IncFIB/IncHI1B plasmids, respectively. The plasmid sizes varied significantly, ranging from 46.1 kb to 307.8 kb; IncC plasmids ranged from 81.2 kb to 307.8 kb, IncX3 was 46 kb, and IncFIB/IncHI1B was 307.8 kb. These plasmids carried ARGs against various clinically relevant antibiotics (β-lactams, aminoglycosides, and fluoroquinolones) (Table 2). There were distinct MGEs, notably Tn*3*, IS*Aba125*, IS*26*, IS*5*, IS*6*, and IS*Kpn14*. Among the four IncC plasmids encoding *bla*_NDM-1_ (pKPP-1, pKPP-2, pKPP-4, and pKPP-5), pKPP-1 and pKPP-4 were highly similar (81.2 kb) but were hosted by different STs (ST2856/KL2 and ST11/KL2), while pKPP-2 and pKPP-5 carried large 307.8 kb and 137.6 kb plasmids, respectively (Table 2).

**Table 2.**
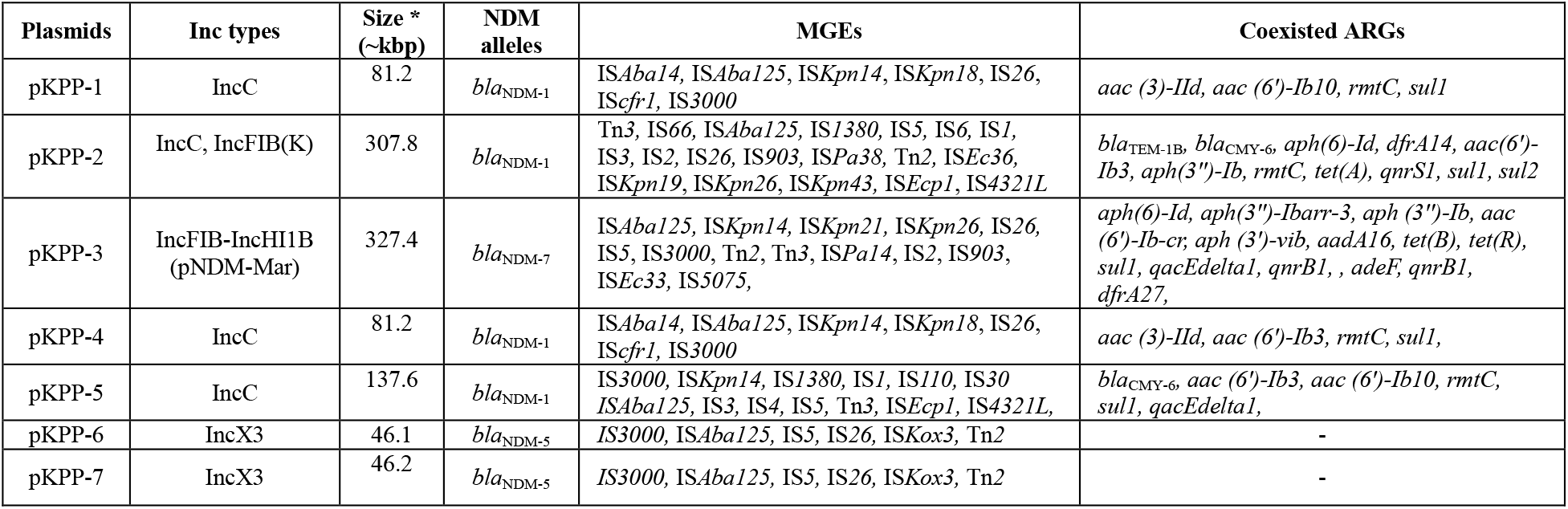
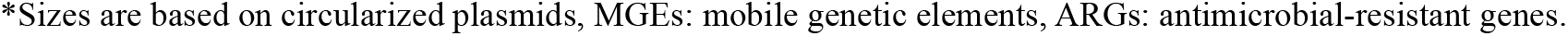
Description of plasmids, antimicrobial resistant determinants, and mobile genetic elements.

^*^Sizes are based on circularized plasmids, MGEs: mobile genetic elements, ARGs: antimicrobial-resistant genes.

The 81.2kb-NDM-1-IncC (pKPP-1 and pKPP-4) plasmid harbored, besides *bla*_NDM-1_ additional genes associated with resistance to sulfonamides (*sul*1), aminoglycosides (*aac(3)-IId, aac(6’)-Ib3*, and *rmtC*). Additionally, the *qac*E gene, which confers resistance to quaternary ammonium compounds such as cetylpyridinium chloride, chlorhexidine, and benzalkonium chloride, was carried by both plasmids. BLASTn analysis revealed that pKPP-1 and pKPP-4 IncC plasmids have high similarity with pKP11ND165-1 (Accession no. CP098372.1) of *K. pneumoniae* from Vietnam with identity and coverage of 100% and 98%, respectively (Figure 2A). The 307kb-NDM-1-IncC (pKPP-2) plasmid coharbored *bla*_NDM-1_ and cephalosporin-resistant genes (*bla*_TEM-1B_, *bla*_CMY-6_), aminoglycosides (*aph (6)-Id, dfrA14, aac(6’)-Ib3, aph(3’)-Ib, rmtC*), sulfonamide (*sul1, sul2*), and tetracycline *(tetA, qnrS1)*. Further, pKKP-5 harbored cephalosporinase gene (*bla*_CMY-6_), aminoglycosides (*aac (6’)-Ib3, rmtC*), and sulfonamide (*sul1*) and showed 100% identity, using BLASTn analysis, with a plasmid (Accession No. CP050164.1) isolated from *K. pneumoniae* in Hong Kong. The IncX3-NDM-5 plasmids (pKPP-6, pKPP-7) carried only *bla*_NDM-5_ and no other ARGs (Figure 2B). The BLASTn analysis revealed 100% identity with pNDM5-SCNJ1 (Accession No. MK715437.1) from China. The IncFIB/IncHI1B-NDM-7-plasmid from pKPP-3, coexisted with an aminoglycoside (*aph(3’’)-Ib, aac(6’)-Ib-cr, aph(3’)-Vib, aadA16, aph(6)-Id*), tetracycline (*tet*B), sulfonamide (*sul*1), fluoroquinolones (*qnrB1*), trimethoprim (*dfrA27*), rifampicin (*ARR-3*) (Figure 2C). Similarity searches against the GenBank database showed partial identity with *bla*_NDM-7_ containing plasmid (pKJNM10C3.2) in *K. pneumoniae*. This plasmid was isolated from a nasal swab of a premature newborn from India (Accession no. NZ_CP030878.1).

**Figure 2.**
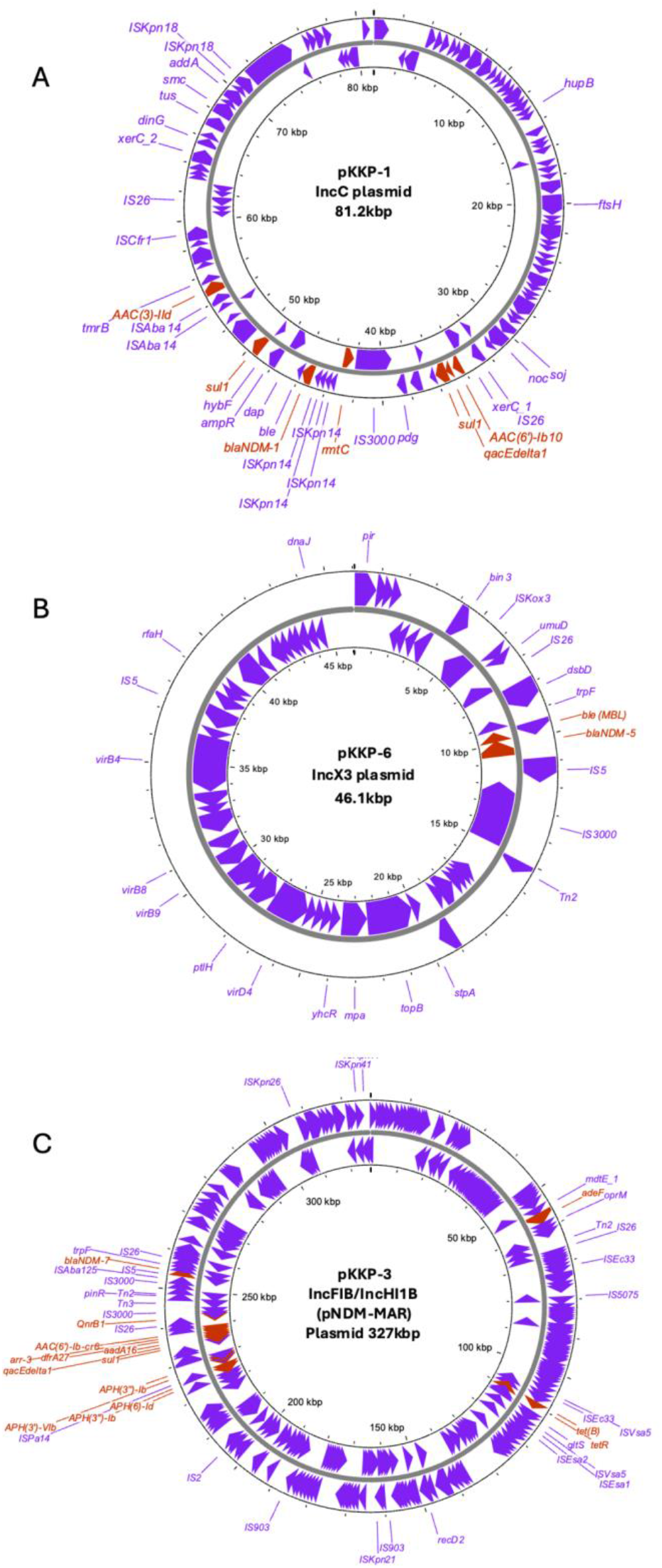
Schematic representation of NDM-harboring plasmids in *K. pneumoniae*. The map of (A) pKPP-1-IncC plasmid harboring *bla*_NDM-1_, (B) pKPP-6 IncX3 plasmid harboring *bla*_NDM-5_ (C) pKPP-3-hybrid IncFIB/IncHI1B plasmid harboring *bla*_NDM-7_. Red arrows in the inner and external rings depict the ARGs, while purple arrows show the various ISs and functional proteins.

### Genetic context of *bla*_NDM_

IS*3000* and IS*Aba14* surrounded the flanking region *bla*_NDM-1_ of pKPP-1 and pKPP-4 plasmids. Insertions of four IS*Kpn14*, IS*3000*, and IS*Aba125* were found upstream of *bla*_NDM-1_, while *ble*_MBL_, *dap, ampR, hybf*, and *sul1* were detected downstream (Figure 3A). The *bla*_NDM-1_ immediate genetic environment showed 93% similarity to a plasmid from *Enterobacter hormaechei* (Accession no. CP115151.1) except for four IS*Kpn14* insertions upstream (Figure 3B). The IS*Kpn14* and IS*Aba125* genes were located upstream of the *bla*_NDM-1_ in the pKPP-2 and pKPP-5 plasmids, while *ble*_MBL_, *trpF, dsbD, cutA, groS, gro*L, and *wapA* were located downstream (Figure 3A). The genomic background shared 100% homology with the IncC plasmid of *K. pneumoniae* (Accession no. CP050164.1) (Figure 3B). The IS*3000* and IS*26* surrounded the *bla*_NDM-5_ genetic structure in the pKPP-6 and pKPP-7 plasmids. Upstream of the *bla*_NDM-5,_ IS*3000*, and IS*Aba125* with insertion of IS*5* were found; however, *ble*_MBL_, *trpF*, and *dsbD* were located downstream (Figure 3A). The genetic context revealed high similarity with pKO_4-NDM-5 of *K. pneumoniae* (Accession no. CP091474.2) (Figure 3B). Furthermore, the genetic structure of pKPP-3 harboring *bla*_NDM-7_ was surrounded by IS*3000* and IS*26*. IS*Aba125* was inserted between the IS*5* and IS*3000* upstream of *bla*_NDM-7_, while the *ble*_MBL_ and *trpf* were located downstream (Figure 3A). This showed a high similarity with pVA04-46 of *K. pneumoniae* (Accession no. CP093504.1), with the difference in IS*3000* size (Figure 3B).

**Figure 3.**
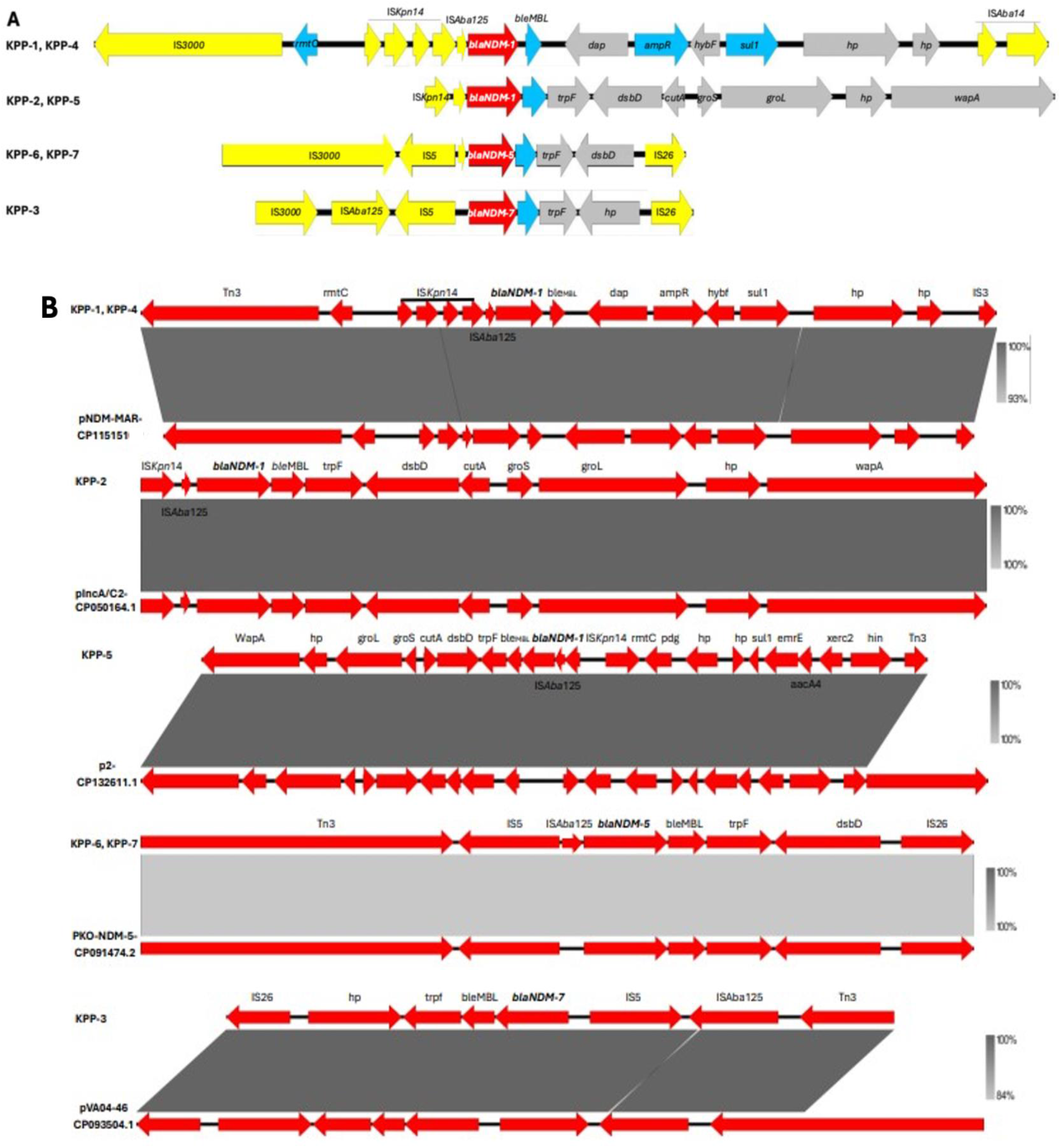
Comparative analysis of the genetic context of *bla*_NDM_ alleles. Figure (A) displays the genetic context with *bla*_NDM-1, -5, -7_ alleles in red arrows, additional ARGs in blue arrows, ISs in yellow arrows, and proteins in gray. The sequences KPP-1, KPP-4, KPP-2, KPP-5, KPP-6, and KPP-7 are all identical. Figure (B) presents the comparative analysis of the genetic context of *bla*_NDM-1, -5, -7_ alleles with previously published sequences (CP115151.1), (CP050164.1), (CP132611.1), (CP091474.2), and (CP093504.1).

## DISCUSSION

Antimicrobial resistance is considered a “silent pandemic” and a public health concern worldwide, particularly in LMICs such as Pakistan (21). This study sheds light on the genetic characteristics of CR-KP isolated from a clinical setting in this country. The present study analyzed CR-KP from blood, pus, and tracheal aspirates belonging to different STs. We found five STs; ST11, ST716, ST16, ST464, and ST2856 in the present study. Carbapenemase-producing ST11 is a well-established high-risk lineage worldwide. A previous study in Pakistan found that CR-KP belonging to ST11 was significantly associated with blood, pus, and urine samples in a clinical setting (20). A One-Health study discovered a high prevalence of *K. pneumoniae* ST11 isolates from human clinical samples and animals (22). Similar ST11 strains were found in a comprehensive arthropod study in South Asia (23). Further, a large-scale study reported *K. pneumoniae* belonged to ST11 in neonatal rectal swabs in Pakistan (24). To our knowledge, *K. pneumoniae* ST716, ST16, ST464, and ST2856 have not been reported in Pakistan; however, a Brazilian study highlighted the emerging high-risk ST16 clone (2). In addition, Asian studies from India and other Asian countries have reported similar STs (ST716, ST16, ST464) (25-28).

Our study on CR-KP revealed a disturbing trend of resistance to most of the “Access” and “Watch” classes of antibiotics, with sensitivity only to “Reserve” classes like colistin, similar to previous reports (24, 29-31). The emergence and spread of AMR are intricately linked with the acquisition of ARGs encoded on various plasmids. Our study identified three *bla*_NDM_ alleles, namely *bla*_NDM-1_, *bla*_NDM-5_, and *bla*_NDM-7_ in CR-KP, mirroring the widespread prevalence of these NDM alleles in other studies (16, 20, 32). However, the genetic context of NDM-7 harbored plasmid has not been reported from Pakistan so far. The interplay of bacterial plasmids and MGEs plays a crucial role in transferring ARGs among bacteria, impacting the prevalence of AMR infections locally and nationally (15). Plasmid replicon typing in this study identified various replicon types, including IncC, IncX3, and the hybrid IncFIB-IncHI1B (pNDM-Mar). The *bla*_NDM-1_ was found to be carried by IncC, *bla*_NDM-5_ by IncX3, and *bla*_NDM-7_ by the hybrid IncFIB-IncHI1B. These plasmids have acquired additional ARGs (*bla*_OXA_, *bla*_TEM,_ *aac*, and *sul1*) and MGEs (Tn*3*, IS*6*, IS*30*, and IS*Aba125*) and are responsible for the dissemination of ARGs in clinical settings. Recently, *bla*_NDM-5_ alleles harbored IncX3 plasmid were identified in the clinical strains of *E. coli* in Pakistan (33). It was observed that over the past decade, there has been a global increase in the prevalence of IncX3 plasmids carrying *bla*_NDM-5_ (34). Previous data also suggested that *bla*_NDM_ is carried by IncC plasmids (18, 35) and *bla*_NDM-5_ harbored (36-38). However, *bla*_NDM-7_ has been identified primarily on IncX3 plasmids (35, 39). To our knowledge, this is the first study to report the presence of *bla*_NDM-7_ carried by IncFIB-IncHI1B plasmids in CR-KP clinical isolates from Pakistan.

In conclusion, we have identified a broad host range of plasmids carrying NDM-1, -5, and -7 alleles in *K. pneumoniae* belonged to ST11, ST716, ST16, ST464, and ST2856. IncC and IncX3 plasmids harbored *bla*_NDM-1_ and *bla*_NDM-5_ respectively, leading to decreased sensitivity to multiple antibiotics, limiting treatment options, and necessitating additional infection control measures. Notably, we have observed the presence of *bla*_NDM-7_ along with other ARGs on a hybrid IncFIB/IncHI1B plasmid in *K. pneumoniae* in Pakistan for the first time. Implementing Pakistan’s National Action Plan on AMR is essential to mitigate its burden. Furthermore, adopting active and effective infection control and prevention measures is crucial to prevent such infections in hospitals.

## MATERIALS AND METHODS

### Bacterial strain identification

The study involved the collection of seven strains of CR-KP from inpatients in a clinical setting in Lahore, Pakistan. These strains were collected from clinical samples in a six-month surveillance study (April to September 2023) for carbapenem-resistant bacteria. Bacterial isolates were confirmed using a Vitek 2 compact system (bioMérieux, France) and a MALDI-TOF mass spectrophotometer (Bruker, USA), following the manufacturer’s instructions.

### Phenotypic antimicrobial susceptibility testing

The Vitek 2 compact system was utilized with adherence to the guidelines of the Clinical and Laboratory Standards Institute (CLSI-M100-S27) to detect the antimicrobial susceptibility of various commonly employed antibiotics, including amoxicillin/clavulanate, ampicillin/sulbactam, ceftriaxone, ceftazidime, cefepime, ciprofloxacin, levofloxacin, co-trimoxazole, amikacin, tigecycline, and colistin. Further, colistin susceptibility was determined by the micro broth dilution assay. Interpretation of the antimicrobial susceptibility results was carried out by the CLSI guideline (40).

### Whole Genome Sequencing

For each isolate, a single bacterial colony was inoculated in 3 mL of LB broth, and 1.5 mL was collected during the exponential phase. Tubes were centrifuged, and the bacterial pellet was washed with 1X PBS. DNA was extracted using the automatic Kingfisher Flex instrument (Thermo Scientific, USA). DNA was quantified using a Qubit fluorometer (ThermoFisher Scientific, USA). 400 ng of high molecular weight DNA were used to prepare sequencing by ligation libraries using the Native Barcoding Kit 24 V14 (SQK-NBD114.24, ONT) that was run in R10.4.1 flow cell in a MinION device (ONT).

### *In silico* analysis

Raw pod5 data was base called on the high computing performance cluster (HPC) Geneva using Dorado (v0.5.1) with the super accuracy model. The resulting fastq files were used for de novo assembly using Flye v2.9.1 and polished with Medaka v1.7.0. Assembled genome sequences were annotated using Prokka (v1.14.5). Sequence types (STs), ARGs, and plasmid types were determined using the Centre for Genomic Epidemiology (v2.0.0) (http://www.genomicepidemiology.org/) web tools such as MLST, ResFinder, and PlasmidFinder. Insertion elements (IS) were annotated using online databases IS Finder (v.27) (https://isfinder.biotoul.fr/). The presence of virulence genes was determined with Virulence Finder (https://cge.cbs.dtu.dk/services/VirulenceFinder/) and the virulence factor database (VFDB, http://www.mgc.ac.cn/VFs/main.htm). Capsule (K) and surface (O) loci of *K. pneumoniae* were determined by Kaptive (v0.7.3) (https://kaptive-web.erc.monash.edu/). Proksee (https://proksee.ca/) was used for the annotation and visualization of plasmids. Easyfig 2.1 was used to align and visualize plasmid alignments.

## Data availability

## Acknowledgements

The author acknowledges support from the Federal Commission for Scholarships for Foreign Students for the Swiss Government Excellence Scholarship (ESKAS No. 2023.0575) for the academic year 2023-24. We thank the sequencing platform of the University of Geneva and Foundation Ernst et Lucie Schmidheiny. We are grateful for the support of the Institute of Microbiology, Faculty of Life Sciences, Government College University, Faisalabad, Pakistan.

